# Physical mechanisms of amyloid nucleation on fluid membranes

**DOI:** 10.1101/2019.12.22.886267

**Authors:** Johannes Krausser, Tuomas P. J. Knowles, Anđela Šarić

**Affiliations:** Department of Physics and Astronomy, Institute for the Physics of Living Systems, University College London, London WC1E 6BT, UK; MRC Laboratory for Molecular Cell Biology, University College London, London WC1E 6BT, UK; Department of Chemistry, University of Cambridge, Lensfield Road, Cambridge CB2 1EW, UK; Cavendish Laboratory, University of Cambridge, Cambridge CB3 0HE, UK

## Abstract

Biological membranes can dramatically accelerate the aggregation of normally soluble protein molecules into amyloid fibrils and alter the fibril morphologies, yet the molecular mechanisms through which this accelerated nucleation takes place are not yet understood. Here, we develop a coarse-grained model to systematically explore the effect that the structural properties of the lipid membrane and the nature of protein-membrane interactions have on the nucleation rates of amyloid fibrils. We identify two physically distinct nucleation pathways and quantify how the membrane fluidity and protein-membrane affinity control the relative importance of those molecular pathways. We find that the membrane’s susceptibility to reshaping and being incorporated into the fibrillar aggregates is a key determinant of its ability to promote protein aggregation. We then characterise the rates and the free energy profile associated to this heterogeneous nucleation process in which the surface itself participates in the aggregate structure. Finally, we compare quantitatively our data to experiments on membrane-catalysed amyloid aggregation of *α*-synuclein, a protein implicated in Parkinson’s disease that predominately nucleates on membranes. More generally, our results provide a framework for understanding macromolecular aggregation on lipid membranes in a broad biological and biotechnological context.

## INTRODUCTION

The aggregation of normally soluble proteins into *β*-sheet rich amyloid fibrils is a common form of protein assembly that has broad implications across biomedical and biotechnological sciences, in contexts as diverse as the molecular origins of neurodegenerative disorders to the production of functional materials [1, 2]. The presence of surfaces and interfaces can strongly influence amyloid aggregation, either catalysing or inhibiting it, depending on the nature of the surface. This effect has previously been studied for the cases of amyloid nucleation on nanoparticles [3–5], flat surfaces [6, 7], and on the surface of amyloid fibrils themselves [8, 9].

Lipid bilayers are a unique type of surface, which is ubiquitous in biology and is the main contributor to the large surface-to-volume ratio characteristic of biological systems. They are highly dynamic self-assembled structures that can induce structural changes in the proteins bound to them [10, 11] and markedly affect protein aggregation propensities [12, 13]. While nucleation on the surfaces of lipid membranes can influence fibril formation dramatically, below the critical micelle concentration alternative surfactant-driven fibrillation pathways in solution have been proposed [14].

Increasing experimental evidence supports the principle that the interaction between amyloidogenic proteins and the lipid cell membrane catalyses *in vivo* amyloid nucleation, which is involved in debilitating pathologies. Remarkably, through surface-driven catalysis lipid bilayers can enhance the kinetics of *α*-synuclein aggregation, the protein involved in Parkinson’s disease, by over three orders of magnitude with respect to nucleation in solution [15].

Bilayer membranes can exist in different structural phases and can undergo local and global phase changes. A large body of work has focussed on exploring how the membrane’s dynamical properties, such as its fluidity, relate to amyloid aggregation of bound proteins [16–23].

For instance, fluid membranes, constituted of short and saturated lipid chains, were found to most effectively catalyse the nucleation of *α*-synuclein [16], while less fluid membranes composed of long lipid chains had less catalytic power. Furthermore, the addition of cholesterol to lipid membranes was found to alter its fluidity and govern the nucleation rate of A*β*42 [22], a peptide implicated in Alzheimer’s disease. In these cases the physical properties of the membrane are controlled through variations in its composition, and decoupling the role of the membrane’s physical properties from its chemical specificity is extremely challenging.

The questions we focus on here is how the microscopic steps that drive amyloid nucleation at the membrane surface are altered by the inherently dynamic nature of lipid bilayers.

Computer simulations can be of great help in this case, enabling us to systematically investigate the role of the physical and chemical properties of lipid membranes independently from one another, thus helping to identify key players behind membrane-driven amyloid nucleation.

In this work, we develop a coarse-grained Monte-Carlo model for studying the nucleation of amyloidogenic proteins on lipid membranes. We use it to identify the microscopic mechanisms which connect the membrane fluidity, the rate of amyloid nucleation, and the morphology of amyloid aggregates. We find that the membrane most efficiently catalyses amyloid nucleation by donating its lipids to the nucleating fibril, which depends *i*) on the lipid solubility and often correlates with membrane fluidity and *ii*) the affinity of proteins to the membrane. This interdependence controls both the morphology of the resulting aggregates, which can range from protein-rich to lipid-rich, and the rate of fibril formation. We then discuss how our results provide a mechanistic explanation for a number of recent experimental observations and offer a platform for studying strategies for bypassing amyloid nucleation in a cellular context.

## RESULTS

### Computational model

To study the essential features of membrane-assisted nucleation, we develop a coarse-grained computational model that takes into account the dynamic nature of the lipid membrane, the process of membrane-bound protein oligomerisation, and the protein’s structural transition that allows fibril formation.

The lipid bilayer membrane is described using a previously published three-beads-per-lipid model [24], where the two hydrophobic tail beads are mutually attractive, allowing for the formation of a stable bilayer. We control the membrane’s thermodynamic phase state by varying the depth of the interaction potential between the lipid tails, *k*_B_*T*/*ε*, where *k*_B_ is the Boltzmann constant and *T* is the temperature. In particular, to evaluate the protein nucleation kinetics across different membrane phases, we tune *k*_B_*T*/*ε* between 0.775 (gel phase) and 1.135 (fluid phase), as shown in Fig. 1(c) (see Methods for details).

**FIG. 1.**
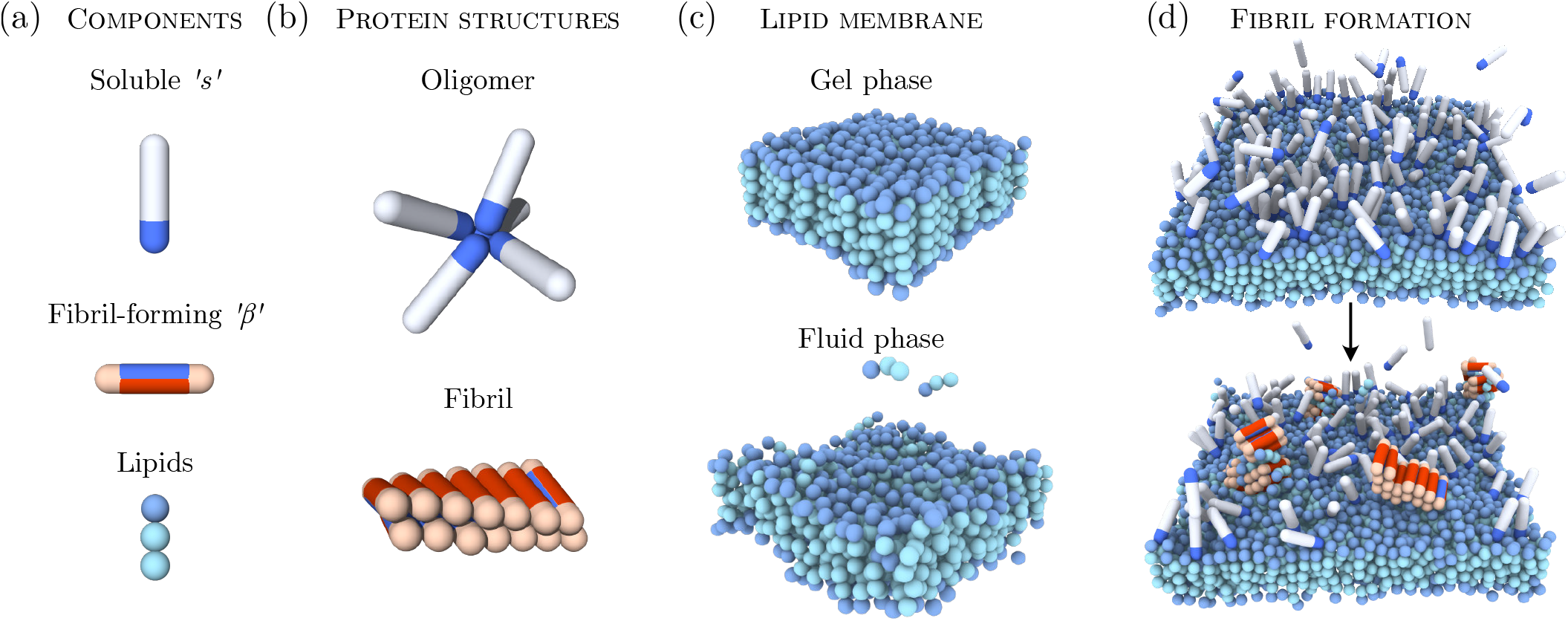
Simulation model and membrane-driven fibril formation. **(a)** Proteins can exist in two distinct conformations: a soluble ‘*s*’ and a *β*-sheet-prone conformation. Lipid molecules are modeled by one hydrophilic head and two hydrophobic tail beads. **(b)** Soluble proteins can form oligomers via their tip-to-tip interactions. Protein molecules in the *β*-sheet-forming conformation can assemble into fibrillar structures through the interactions of the blue side-patches. **(c)** The lipid membrane can exist in different structural phases depending on the inter-lipid interactions set in the model. Shown here are the gel phase (*k*_B_*T*/*ε* = 0.775) and fluid phase (*k*_B_*T*/*ε* = 1.135). **(d)** Example showing the formation of an amyloid fibril on the lipid membrane from bound proteins.

The minimal model for the amyloid-forming proteins is based on the coarse-grained model introduced in Refs. [25, 26]. Proteins are modelled as hard patchy spherocylinders and can exist in two distinct conformational states: a soluble (*s*) and a *β*-sheet forming (*β*) conformation, both of which are equipped with different arrangements of interaction patches. This model captures the aggregation behaviour through the colloidal nature of the building blocks, and the internal dynamics of the protein through the internal degree of freedom associated to the *s*-to-*β*-sheet conformational transition.

Soluble proteins interact through an attractive end cap on the spherocylinder, which allows for the formation of unstructured protein oligomers, Fig. 1(a). The *β*-prone conformation has an attractive side patch instead of a cap, which mediates attractive interactions both with the *s* and *β* conformations facilitating the alignment of proteins and the formation of elongated fibrils. The *s*-state represents the conformational ensemble of protein molecules in their soluble states, for instance a random coil for both the *Aβ* peptide and *α*-synuclein. The *β*-prone-state describes the a conformation of the polypeptide chain which possesses strong intermolecular interactions as found in the *β*-sheet-rich cores of amyloid fibrils. Therefore, the two spherocylinder states represent different classes of membrane binding modes rather than reflecting precise conformational geometries, which vary for different amyloidogenic proteins.

Crucially, the proteins can undergo a structural transition between the soluble and *β*-sheet-forming state. This stochastic transition is penalised with a free energy barrier Δ*F*_s→*β*_, reflecting the fact that amyloidogenic proteins lose conformational entropy when converting from native to the *β*-sheet-prone state and are rarely found in the *β*-sheet conformation on their own.

Analysis of the simulation trajectories shows that membrane binding is first achieved by the soluble protein conformation, whose affinity for the lipid membrane heads is set by the value of *ε*_sm_. To model the amphipathic nature of the protein and allow it to partially insert and anchor into the lipid membrane, as observed for both *α*-synuclein [27] and A*β* [28], the protein can also interact with the lipids tails with a fraction of *ε*_*s*m_. The *β*-sheet-prone conformation is equipped with two separate patches on opposing sides of the spherocylinder with affinity for either the proteins or the lipids, Fig. 1(a). Such a twofold binding motif enables the protein to form a fibril and, at the same time, interact with the lipid membrane. This motif is chosen to mimic the general membrane-binding characteristics of amphipathic proteins such as *α*-synuclein and *Aβ*, which upon binding to the membrane can undergo a structural transition that promotes *β*-sheet formation in contact with lipids [10, 29, 30]. The strength of the lipophilic attraction of the protein (red patch) is controlled by *ε*_*β*m_, while the protein-protein interactions of the *s*- and *β*-states (blue patches) are controlled by *ε_ss_*, *ε_sβ_* and *ε_ββ_*.

Using the model presented in Ref. [25] we found that no nucleation occurs in solution during the simulation time, while nucleation in the presence of lipids can be fast, as observed in experiments [31]. To be able to explore the mechanisms of membrane-assisted nucleation across different conditions in a computationally efficient way we set the conversion barrier Δ*F*_s→*β*_ and the *β*-membrane interaction *ε*_*β*m_ proportionally to 10 *k*_B_*T* while suppressing nucleation in solution. Further details can be found in the *Supporting Information*.

### Membrane-assisted nucleation mechanisms

Through binding to the bilayer, the soluble protein molecules readily arrange into small unstructured oligomers on the membrane due to their self-interaction. The oligomerisation on the membrane is more efficient than in solution due to the higher protein concentration on surface. If after a lag time a successful conversion in one of the proteins to its *β*-prone conformation has occurred, aggregation and potential fibril elongation can start. Our simulations show that the properties of lipid membranes can dramatically influence the pathway through which this nucleation occurs. In the following, we distinguish two basic pathways and the related morphologies of amyloid aggregates: a protein-rich and a lipid-rich, which also give rise to an intermediate case, as illustrated in the phase diagram in Fig. 2.

**FIG. 2.**
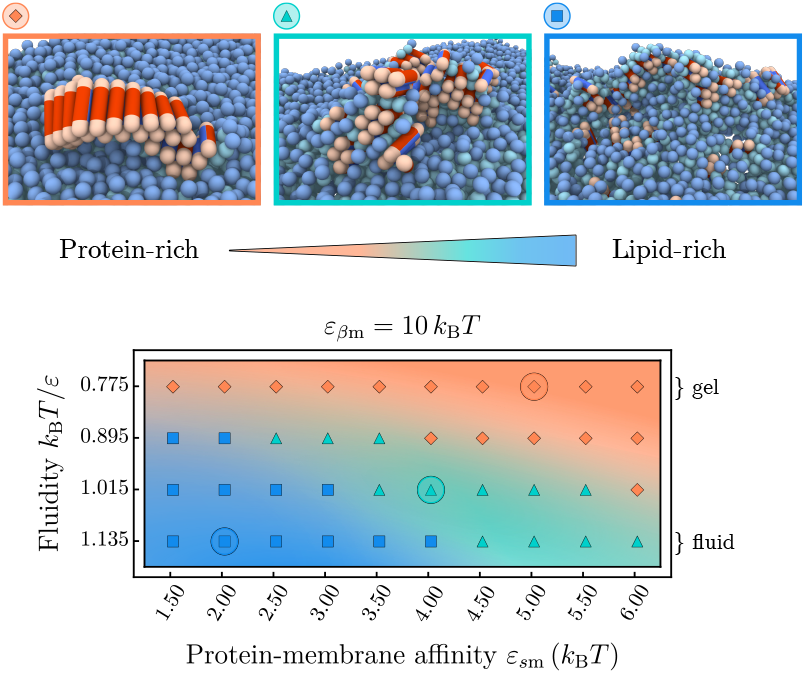
Morphologies of protein-lipid aggregates. Phase space of the protein-lipid cluster morphologies depending on the membrane fluidity and protein-membrane affinity. Three main areas can be distinguished: extended fibrils (orange), smaller fibrillar clusters with interstitial lipids (green), and strongly mixed lipid-protein clusters (blue). The representative snapshots correspond to the circled parameter values; soluble proteins are not shown.

In the gel phase (*k*_B_*T*/*ε* = 0.775), membrane lipids are packed closely and bound proteins are unlikely to penetrate into the bilayer. In this case, the membrane essentially behaves as a static surface: proteins adsorb onto the surface, form transient oligomers, which eventually provide an environment for fibril nucleation [25]. Such a pathway typically results in the appearance of elongated fibrils epitaxially growing on top of the bilayer (orange area in Fig. 2), which often detach and diffuse away from the membrane. The size of the fibrils strongly depends on the protein-membrane affinity ε_sm_. High ε_sm_-values lead to an increased membrane coverage, which causes fast growth of the fibril after initial conversion, and hence a higher proportion of longer fibrils, as shown in Supplementary Fig. A5.

The morphology of nucleated clusters changes distinctly when the membrane is in the fluid phase, as best seen in the regime of low protein-lipid affinities in Fig. 2. In this case lipids are comparatively weakly bound within the bilayer and can be extracted from it more easily. In fact, experiments show that the lipid solubility is increased when shortening the acyl chain length of saturated lipids [16] (see *Supporting Information* for details. This enables the lipids to actively participate in the formation of fibril nuclei. At low protein-lipid affinities *ε*_*s*m_ the membrane is weakly covered by proteins and the first nucleation step typically proceeds via the direct interaction of a single *β*-prone protein with the hydrophobic tail of a lipid, either by lipid extraction from the bilayer or partial insertion of the protein into a packing defect. The converted *β*-prone protein can then get coated in lipids or (further) inserted into the lipid bilayer, which hampers the fast elongation of fibrils and leads to mixed protein-lipid clusters (blue area in Fig. 2). This nucleation pathway is promoted by packing defects in the lipid bilayer at increased membrane fluidities. The exact composition of the aggregates depends on the relative rate of incorporation of lipids and proteins into an aggregate, governed both by the membrane fluidity and the protein-membrane affinities.

At higher protein-membrane affinities 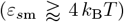 the membrane is substantially covered by proteins, see Supplementary Fig. A2. The local environment of a growth-competent nucleus will hence be abundant in soluble proteins, leading to a faster addition of protein monomers and hence fibrils with lower lipid content (green area in Fig. 2). Interestingly, the bound proteins also modify the local phase state of the membrane, as indicated by a reduction of the average area per lipid as ε_sm_ is increased (Supplementary Fig. A4). This hinders lipid extraction and yields an intermediate nucleation pathway between the lipid-rich and protein-rich regimes, as illustrated in Fig. 2.

Similar trends to those observed in our simulations have been reported in atomic force microscopy experiments monitoring A*β* aggregating on model lipid membranes [20]. On fluid model membranes with strong electrostatic interactions between proteins and lipids, bilayer deformations and clustering of lipids around A*β* were observed, whereas on gel phase membranes elongated mature fibrils appeared, just like in our simulations. Furthermore, experiments have shown that *α*-synuclein fibril formation in the presence of vesicles leads to membrane remodelling and lipid extraction [32], yielding fibrils intercalated with lipids that often cause vesicle disintegration. According behaviour involving membrane rupture can be observed in our simulations at later stages of the aggregation process when stresses induced by the growing fibril-lipid aggregates become too large.

### Membrane fluidity enhances nucleation rates

In addition to controlling the morphologies of amyloid aggregates, the membrane has an immediate effect on the rates of amyloid nucleation. To convert from the soluble into the fibril-forming state the protein needs to overcome the intrinsic free energy barrier Δ*F*_*s*→*β*_. The role of the membrane in modifying this nucleation barrier can be twofold: (*i*) to increase the local concentration of proteins by restricting their mobility to the membrane surface and (*ii*) to actively participate in the formation of the pre-fibrillar nucleus through hydrophobic interactions. Here, we decouple these two effects by analysing two separate scenarios: a control case in which a protein is only allowed to adsorb onto the membrane but cannot co-nucleate with lipids, and another one in which *β*-sheet-prone proteins can co-nucleate with lipids as depicted in Fig. 2. The two scenarios are characterised by the presence or absence of the lipid-protein interaction patch of the *β*-prone conformation (red patch), while the interaction between the *s*-conformation and the lipids remains unaltered.

Starting with the control case, we remove the red patch by setting *ε*_*β*m_ = 0 *k*_B_*T*. As evident from Fig. 3(a), the nucleation rate *r*(*ε*_*s*m_) is a non-monotonic function of the protein-membrane affinity *ε*_*s*m_. Here, the rate *r* is the inverse mean lag time of *β*-prone protein dimer formation. At small *ε*_*s*m_-values the nucleation rates are low across all fluidities, where virtually no proteins are adsorbed onto the membrane. At intermediate membrane-protein affinities, the formation of stabilised membranebound oligomers lowers the nucleation barrier and increases the nucleation rates. At high membrane-protein affinities the nucleation process is inhibited due to the unfavourable free energy associated with nucleus detachment. Indeed at *ε*_*β*m_ = 0 *k*_B_*T* upon conversion to the *β*-state the protein loses its interaction with the membrane, which becomes costly at high protein-membrane affinities *ε*_*s*m_, hence prohibiting nucleation. The onset of this regime is systematically shifted to higher *ε*_*s*m_-vales as the fluidity is increased. This is rooted in the fact that a higher lipid mobility inhibits the formation of stable oligomers on the membrane surface (Supplementary Fig. A3).

**FIG. 3.**
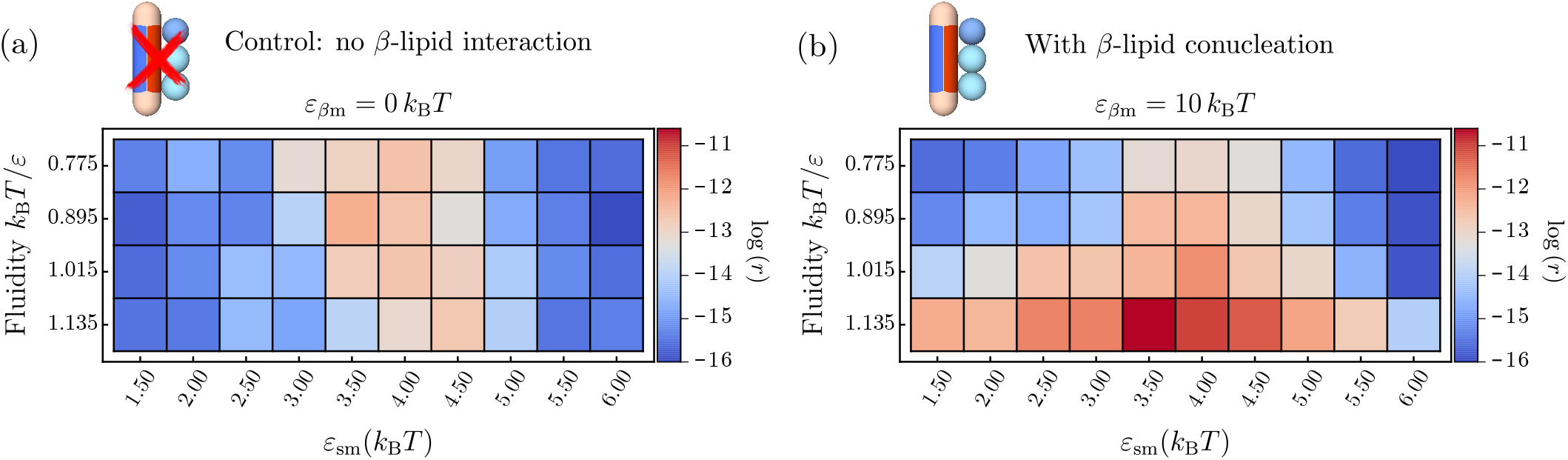
Amyloid nucleation rates on the membrane. **(a)** Control case: nucleation rates when only soluble proteins interact with the lipid beads and *β*-like proteins do not penetrate into the hydrophobic core. Non-monotonic behaviour is observed with respect to the protein-lipid interaction *ε*_*s*m_. **(b)** Nucleation rates when the interactions between the proteins and the lipid-tails are non-zero and proteins can penetrate the bilayer, as in Fig. 2. Amyloid nucleation is sped up in the direction of higher fluidities in addition to the non-monotonic scaling with *ε*_*s*m_ observed in (a).

In the second scenario, the *β*-sheet-prone protein conformation carries a side patch with an affinity for the hydrophobic lipid tails of *ε*_*β*m_ = 10 *k*_B_*T*, rendering the protein amphipathic. The value of *ε*_*β*m_ is chosen larger than *ε*_*s*m_ to reflect the stronger membrane binding associated to a higher *β*-sheet content found in experiments [33]. Strikingly, the presence of the *β*-lipid interaction leads to a drastic change in the nucleation rates at higher membrane fluidities across all parameters pairs investigated, as shown in Fig. 3(b). This effect is caused by the progressive exposure of the membrane’s hydrophobic core and the concomitant hydrophobic contacts between lipids and proteins. Loose lipid packing and the enhanced mobility of the lipids both in and out of the membrane plane enables the participation of lipids in the s-*β* conformational change and the formation of fibrillar clusters.

The inclusion of lipids in protein aggregates efficiently drives fibril nucleation as the interaction between the lipid tails and proteins in the *β*-sheet prone conformation is favourable, which reduces the free energy barrier of the *s* → *β* conversion and stabilises fibril nuclei. This effect is more pronounced at increasing membrane fluidities, which allow for better access to the hydrophobic regions of the lipid bilayer (Fig. 3). At the same time, a sufficiently but not too strong protein-membrane binding affinity *ε*_*s*m_ enables both efficient binding and oligomer formation on the membrane without prohibiting the conformational conversion.

### Free energy barrier for nucleation

To further characterise the molecular mechanisms of membrane-driven catalysis of amyloid aggregation, we investigate the free energy barriers connected to the different nucleation pathways. To this effect we sample the free energy landscape for the *s* → *β* conformational conversion of a protein along the protein-membrane centre-of-mass separation *z_cm_* in different membrane environments, thereby setting aside the effect of surface oligomerisation of the soluble proteins. First, we evaluate the free energy profile associated to a protein in either the s or *β* conformation interacting with the membrane, while separately varying the membrane fluidity and protein-membrane affinity, see Fig. 4.

**FIG. 4.**
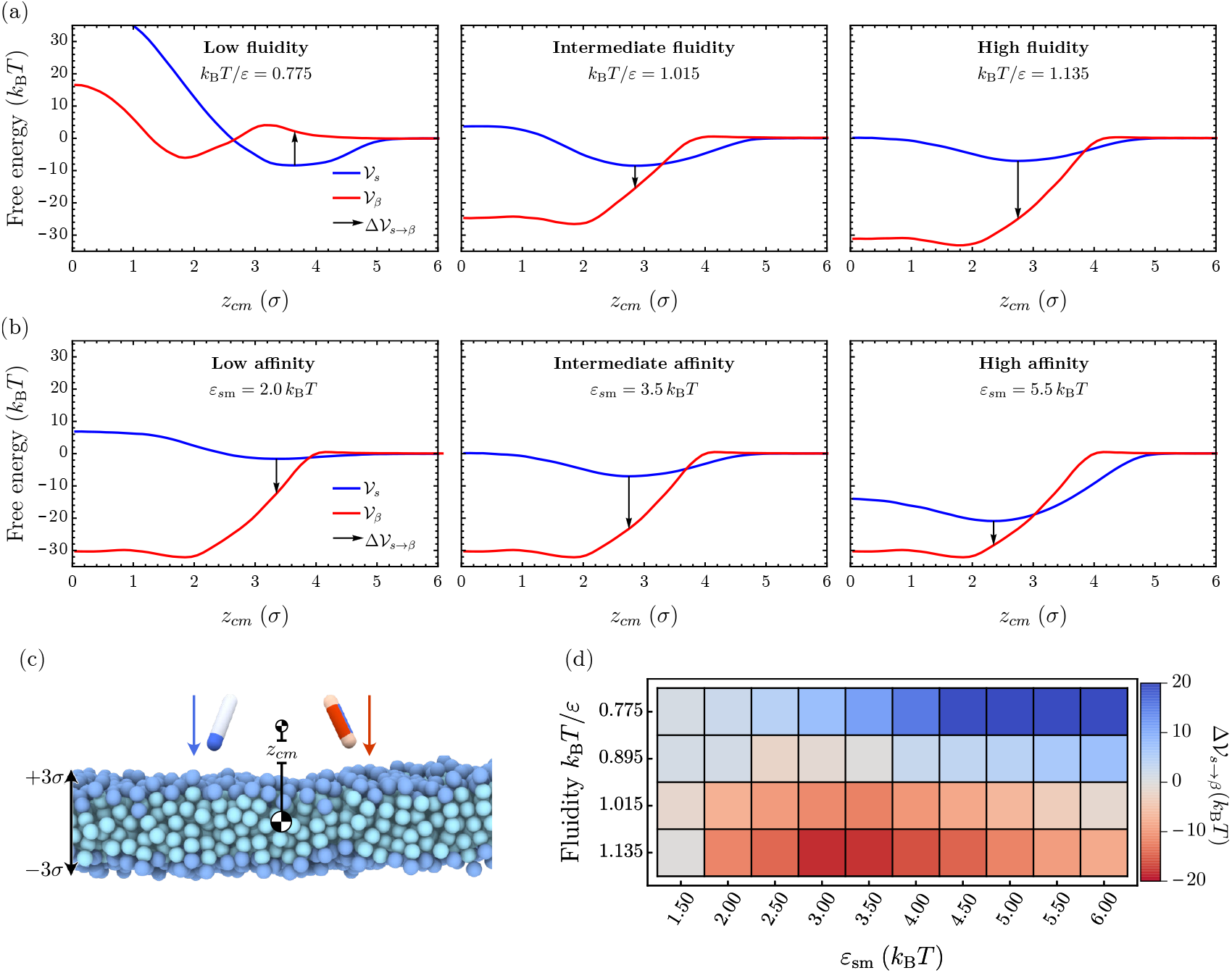
Changes in the potential of mean force at increasing membrane fluidity: **(a), (b)** The three panels each show the free energy profiles 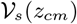 (red) and 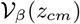 (blue) at increasing fluidities *k*_B_*T*/*ε* = 0.775,1.015, and 1.135 while protein-membrane interactions are kept fixed at *ε*_*s*m_ = 3.5 *k*_B_*T* and *ε*_*β*m_ = 12 *k*_B_*T* **(a)**, and at increasing protein-membrane affinities *ε*_*s*m_ = 2.0,3.5, and 5.5 *k*_B_*T* while the fluidity is kept fixed at *k*_B_*T*/*ε* = 1.135 and *ε*_*β*m_ = 12 kBT **(b)**. The arrows indicate the free energy cost for conformational conversion 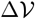, providing a proxy for the nucleation barrier. **(c)** Initial snapshot of umbrella simulations for both particle species. **(d)** Difference between potentials of mean force in the ‘*s*’ and ‘*β*’ conformation evaluated at the minimum of 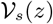 as a function of the membrane fluidity and the membrane-protein affinity *ε*_*s*m_.

Our simulations show that the free energy of binding between the membrane and the protein decreases with increasing fluidity, as evidenced by the results in Fig. 4(a) for *ε*_*s*m_ = 3.5 *k*_B_*T* and *ε*_*β*m_ = 12.0 *k*_B_*T* (see also Supplementary Fig. A8). Concurrently, we observe a softening of the repulsive contribution to the free energy profile 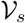 and a shift of its minimum to smaller *z_cm_*-values. This change is the combined result of membrane thinning and a deeper average insertion of the protein into the bilayer at high fluidities. Larger values of the membrane-protein affinity *ε*_*s*m_ at fixed fluidity entail both a deeper free energy minimum and a shift of the minimum position deeper into the hydrophobic core (Fig. 4 and Supplementary Fig. A8). In the case of the protein in the *β*-conformation, the free energy profile 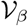 in the gel phase consists of a minimum located inside of the bilayer separated from the membrane surface by a small repulsive barrier, since it is energetically unfavourable for the *β*-protein to insert into the membrane at tight lipid packing (left panel in Fig. 4(a)). Increasing fluidity removes this barrier and 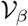 consists of one deep well representing the strong hydrophobic binding of the protein within the lipid membrane.

To estimate the free energy barrier for conversion between the protein’s *s*- and *β*-conformation we evalaute the free energy difference 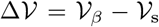 at the equilibrium position of the soluble protein (indicated as a black arrow in Fig. 4). This free energy jump provides a proxy for the nucleation barrier in the dynamic Monte-Carlo simulations where soluble proteins first bind to the lipid and find their equilibrium position before slowly converting into the *β*-sheet-prone conformation. As evident from Fig. 4(a), as the membrane fluidity increases, the nucleation barrier 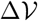 gradually disappears. Deeper insertion of the soluble protein permits strong interactions between the *β*-protein and the hydrophobic core of the membrane upon conformational conversion. Correspondingly, 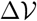 becomes large and negative at high membrane fluidities, resulting in the increase in the nucleation rates (Fig. 4(d)).

Changing the affinity *ε*_*s*m_ can additionally influence the barrier. In the gel phase protein insertion into the membrane is not possible and we observe a monotonic increase of 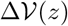 with higher protein-membrane affinities (top row of Fig. 4(d)). In the rate measurements reported in Fig. 3, the nucleation rate initially increased with *ε*_*s*m_ (lower 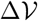) by virtue of surface oligomerisation. This is a multi-protein effect not captured at a level of a single protein conversion. Conversely, 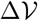 becomes nonmonotonic in *ε*_*s*m_ even without the multi-protein effects at high fluidities, since soluble proteins can partially insert into the fluid membrane. The interaction between the *β*-protein and lipid tails increases closer to the membrane centre, hence the free energy gain of transitioning to the *β* conformation also grows. In the strong binding regime, however, the effective binding free energy of the soluble protein grows faster than that of the *β*-prone protein, leading to the re-entrant increase in the free energy barrier 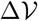 for the conformational conversion.

### Quantitative comparison with experimental data

In what follows, we compare our findings to experimental results [16], which characterise the effect of lipid chemistry on the primary nucleation rates of *α*-synuclein in the presence of lipid membranes.

Considering fully saturated lipids of different acyl chain lengths, the experimental data show that vesicles consisting of the longest acyl chains that form gel-like membranes result in the slowest amyloid aggregation rates. Conversely, the shortest lipid molecules, which have the highest solubility in water and constitute a vesicle in the most fluid phase, lead to the fastest aggregation rates [16].

Since our model is highly coarse-grained and general in nature, it remains non-trivial to map the exact lipid bilayer phase state and protein-membrane interaction parameters from experiments to our model.

Importantly, the lipid model used here does not account for the complex structures of unsaturated acyl chains. The comparison with experimental data is therefore restricted to fully saturated lipids, where membrane fluidity and lipid solubility are directly related.

Nevertheless, the increase in the area per lipid from experiments between the gel phase and the two fluid phase vesicles under consideration is 25 and 31%, respectively. This range closely matches what we observe in simulations and falls in the medium fluidity regime, as shown Fig. 5. This is a parameter-free measurement that we can use to describe the membrane phase state and compare the simulation and experimental results.

**FIG. 5.**
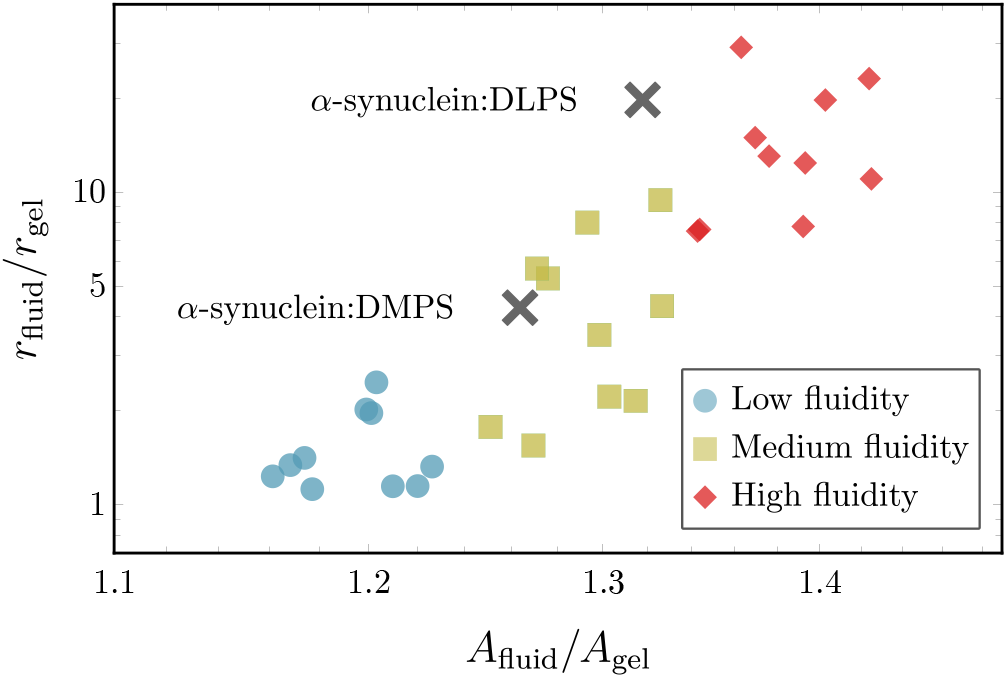
Comparison of simulation results with experimental data: Comparison of the relative increase in nucleation rates *r*_fluid_/*r*_gel_ between the gel phase and fluid membranes as a function of the relative increase in area per lipid *A*_fluid_/*A*_gel_. Coloured symbols show simulation results at different fluidities: (purple) low fluidity (*k*_B_*T*/*ε* = 0.895), (green) medium fluidity (*k*_B_*T*/*ε* = 0.895), and (red) high fluidity (*k*_B_*T*/*ε* = 0.895) with each subset containing 10 different *ε*_*s*m_-values. The two black symbols represent experimental data from Ref. [16]. The experimental values for *r*_fluid_ and *A*_fluid_ of the fluid-phase DMPS and DLPS vesicles are normalised to *r*_gel_ and *A*_gel_ of a gel-phase DPPS vesicle, respectively.

For a quantitative comparison of the nucleation rates, we consider the relationship between the increase in nucleation rates and area per lipid for different fluid-phase membranes relative to the gel phase, i.e. the quantities *r*_fluid_/*r*_gel_ and *A*_fluid_/*A*_gel_.

Considering the scenario in simulations where attractive interactions between the fibril forming protein and hydrophobic core of the bilayer are enabled, i.e *ε*_*β*m_ = 10*k*_B_*T* (Fig. 3 (b)), we observe that enhanced nucleation rates directly correlate with increases in area per lipid, as shown in Fig. 5. Nucleation rates depend sensitively on the membrane structure since direct interactions with lipids tails allows the the fibril-forming proteins to sense packing defects. This is not the case for the non-interacting case (*ε*_*β*m_ = 0*k*_B_*T*), where we find practically no variation of the speed up with changes in the area per lipid, as demonstrated in Fig. A9.

Remarkably, the match between the simulation and experimental data is achieved only if direct interactions between the membrane core and the fibril-forming protein are present. This indicates that the fluidity-dependent nucleation mechanism proposed here is necessary to induce the experimentally observed enhancement in nucleation rates on fluid membranes. Merely surface-assisted nucleation through oligomers without lipid-protein co-nucleation is not sufficient to explain the experimental data.

Hence, the minimal model developed here appears to capture the key physics needed to reproduce and explain the experimental data on membrane-driven amyloid nucleation.

## DISCUSSION

In this work we presented a coarse-grained simulation model to investigate how the phase state of a lipid membrane and membrane-protein binding affinity control the nucleation rates of amyloid fibrils. Depending on the specific amphipathic interaction motif of the amyloidogenic proteins, we identified two main nucleation mechanisms:

1. The formation of protein oligomers on the membrane surface and subsequent nucleation into pure fibrils
2. The mixing of membrane lipids with fibril-forming proteins enhanced by membrane fluidity.

The first mechanism determines the aggregation process when the protein cannot interact with the membrane hydrophobic core. The efficiency of this mechanism is dictated predominantly by the protein-membrane affinity, which stabilises oligomers on the membrane surface. The enhancement of nucleation is limited to a narrow regime of protein-membrane interaction in this case. Interestingly, this effect can be inhibited at high lipid mobilities, where the formation of stable oligomers becomes prohibited. This nicely illustrates that the membrane fluidity *per se* does not catalyse nucleation, but that specific interactions between the protein and the lipids, which often correlate with lipid fluidity, are required.

In fact, when interactions of the protein with the hydrophobic membrane core are present, a second more powerful nucleation mechanism is enabled. In this regime lipids can co-nucleate with proteins, which effectively lowers the nucleation barrier for their conformational conversion and can result in the formation of lipoprotein clusters. This mechanism depends crucially on the membrane fluidity and is enhanced by the presence of packing defects and lipid extraction from the bilayer. Indeed, considering the scenario where the protein in the *β*-sheet-prone state interacts with the lipid tails, we observe a 250-fold speed-up of the nucleation rates between the slowest and fastest cases in the gel and fluid phase, respectively, (Fig. 3b). This is contrasted by only a 20-fold speed-up for the case when the protein cannot interact with the lipid tails (*ε*_*β*m_ = 0 *k*_B_*T*) shown in Fig. 3(a).

Notably, we demonstrate that the window of effective nucleation is significantly broadened by the interaction between the protein and the membrane core leading to even more efficient amyloid nucleation over a wider range of membrane fluidities and protein affinities.

It is important to note that the membrane fluidity in our model is controlled only by inter-lipid interactions, hence the fluidity and the ability of lipids to be extracted from the bilayer are necessarily correlated in our model. This is a good representation for saturated lipids, where lipid solubility controls the phase state of the membrane. However, membrane phase behaviour can also be controlled by lipid geometry, as in the case of polyunsaturated lipids. In such a system the membrane fluidity and lipid solubility are not necessarily correlated in a straightforward way [15]. Similarly, membrane inclusions such as cholesterol [22] or proteins can also control the membrane phase behaviour and have a non-trivial effect on the ability to extract lipids from the bilayer. Our simulations do not capture these more complex couplings.

We found that the nature of the interactions between the protein and the membrane is of key importance in determining the aggregation pathway and the protein’s capacity to disrupt membranes. In our model, the choice of two separate side patches is necessary to capture the incorporation of lipids into amyloid structures observed in experiments. Without a separate protein-lipid patch lipids are pushed out of fibrils and mixed aggregates cannot be achieved ruling out the pathway to lipid-rich aggregates.

Hence, the imbalance between membrane-membrane and membrane-protein interactions is an important factor not only in controlling the rates and pathways of amyloid aggregation, but also in determining whether amyloid aggregates are highly cytotoxic or not. This is also highlighted in a recent experimental study which shows that *α*-synuclein oligomers require a *β*-sheet core to insert themselves into the membrane bilayer and drive disruption of the bilayer [34].

The physical principles identified in this work are general in nature and applicable to a wide range of amyloid-forming proteins, irrespective of their sequence or fold. Even more, the presented mechanisms can also be of relevance to membrane-driven aggregation of other proteins that involve conformational changes and hydrophobic interactions. The computer model developed here also opens the way for testing strategies to bypass membrane-assisted amyloid aggregation, which can for instance involve targeting the interaction between proteins and membrane hydrophobic core, or altering the lipid composition to prevent protein-lipid co-nucleation.

## ACKNOWLEDGMENTS

We are grateful to T. C. T. Michaels for reading the manuscript. We acknowledge support from the Academy of Medical Science (JK and AS), the Cambridge Centre for Misfolding Diseases (TPJK), the BBSRC (TPJK), the Frances and Augustus Newman foundation (TPJK), the ERC grant PhysProt (agreement n° 337969), Wellcome Trust (AS and TPJK), the Royal Society (AS), and the UK Materials and Molecular Modelling Hub for computational resources, which is partially funded by EPSRC (EP/P020194/1).

## METHODS

### Simulation model

The coarse-grained simulation model employed in this study merges a minimal model of amyloid nucleation in solution [25] with a lipid bilayer implicit solvent model [24]. Proteins are represented by hard spherocylinders of diameter *σ* and length *ℓ* = 4*σ* and can exists in a soluble and a *β*-sheet forming conformation, as discussed in the main text. The membrane lipids are represented by a one head bead and two tail beads. Both the dynamics of the membrane-protein system and the conformational switches of the proteins are simulated using a Metropolis Monte-Carlo algorithm.

As depicted in Fig. 1(a), the soluble protein has an attractive end cap which allows it to form oligomeric structures. The pair-wise interaction between two soluble protein tips at distance *r* = |***r***| is given by

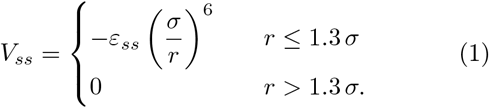

with *ε_ss_* = 4 *k*_B_*T* in this study. In the *β*-sheet forming conformation the spherocylinder has two attractive side patches instead of a tip interaction. One patch with affinity for the membrane, the other with affinity for other proteins.

The attractive protein side patch has the length of 0.7*ℓ* and an opening angle of 180°. If the two patches are facing each other mediates the interaction potential

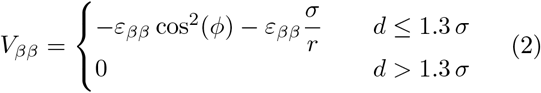

where *d* is the minimal distance between the interacting hydrophobic patches and *ε_ββ_* = 60 *k*_B_*T*.

The interaction between soluble and *β*-prone proteins is set by

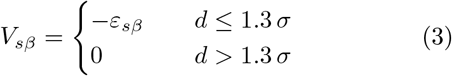

where *d* is the shortest distance between the attractive tip of the soluble protein and the side patch of the *β*-prone protein and *ε_sβ_* = *ε_ss_* + 1*k*_B_*T*.

The coarse-grained implicit solvent model used here for the lipid membrane is defined in Ref. [24]. The membrane consists of three-bead lipids, which self-assemble into a stable bilayer.

The binding of the soluble protein to the membrane lipids is controlled by the interaction potential

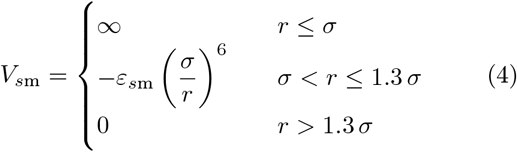

where *ε*_*s*m_ is scaled by 1, 1/2, or 1/4 if the interacting lipid bead is either the head bead, the first or the second tail bead.

The second lipophilic side patch on the *β*-prone protein also has a length of 0.7*ℓ* and an opening angle of 180° and is oriented opposite to the *V_ββ_*-patch. It interacts only with the hydrophobic tail beads of the lipids via

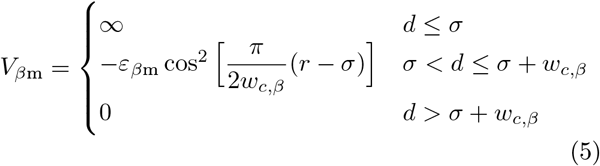

where *d* denotes the minimal distance between the attractive patch and the corresponding lipid bead. The range of *V_βm_* between the proteins and the lipid is set to *w_c,β_* = *σ*. The interaction strength *ε*_*β*m_ is set to 0 or 10 *k*_B_*T*, depending on the specific case under consideration.

Lipid beads interact repulsively via a Weeks-Chandler-Anderson potential given by

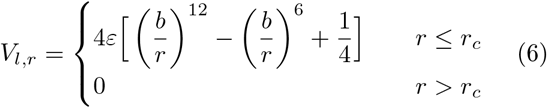

where *r_c_* = 2^1/6^*b*, *b*_head,head_ = *b*_head,tail_ = 0.95*σ*, and *b*_tail,tail_ = *σ*. Beads of a three-bead lipid molecule are connected by two finitely extensible nonlinear elastic (FENE) bonds, described by

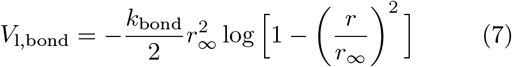

where *k*_bond_ = 30 *ε*/*σ*^2^ and *r*_∞_ = 1.5*σ*. Additionally, the head and second tail bead interact via the bending potential

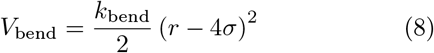

where *k*_bend_ = 10*ε*/*σ*^2^. The hydrophobic interactions between the lipid tails are account for by setting the attractive interaction between the two tail beads to

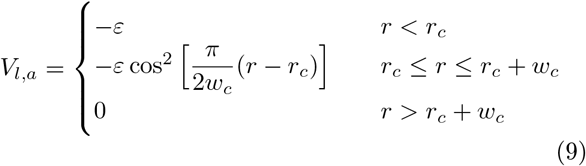

with the *ε* and wc being the depth and the range of the attractive potential, respectively. In this work, we set *w_c_* = 1.42*σ* and vary *ε* to prepare the lipid membrane in different phase state across the gel and fluid regime. In particular, we vary *k*_B_*T*/*ε* between 0.775 (gel phase) and 1.135 (fluid). Note that the gel-fluid phase boundary is approximately located at *k*_B_*T*/*ε* = 0.81.

### Monte-Carlo scheme

The dynamics of the membrane-protein system is simulated by a Monte-Carlo scheme, which includes translational and rotational moves of individual particles. Additionally, the conversion between the soluble and *β*-prone conformation of the protein is also facilitated by a Metropolis criterion. This nucleated conformational change is accepted with probability *p* = min{1, e^−Δ*E*/*k*_B_*T*^}, where Δ*E* denotes the energy difference between the s and *β*-prone state. Switches between the two possible states are attempted with a probability of 0.01 per time step. These conformational changes between the s and the *β* state are penalised by the energy barrier Δ*F*_*s*→*β*_ = 10 *k*_B_*T*.

The simulations were carried out in a cubic simulation box with periodic boundaries in the *x* and *y* directions. The height of the box is kept constant at *L_z_* = 50*σ*. The lengths *L_x_* and *L_y_* are allowed to fluctuate to keep the surface tension of the membrane constant at zero. Soluble proteins are equilibrated according to a grand-canonical ensemble with a fixed chemical potential keeping the concentration of soluble proteins constant in solution. Note that *β*-prone proteins follow the canonical ensemble. The membrane consists of 30 × 30 lipids in each leaf, amounting to 1800 three-bead lipids in total.

In a typical simulation run, proteins initially adsorb to the membrane while being maintained in the soluble conformation. After the equilibrium surface coverage is reached, the proteins are allowed to switch between the soluble and the *β*-sheet-prone state.

## SUPPLEMENTARY INFORMATION

This document includes additional information and results on the simulations carried out in the present study. In particular it includes:

- Overview of interaction in the simulation model
- Remark on nucleation in solution vs membrane-assisted nucleation
- Coverage of the membrane surface by *s*-proteins
- Cluster size distribution of bound *s*-proteins
- Average area per lipid from Voronoi tesselation
- Cluster size distribution of nucleated *β*-proteins
- Dependence of nucleation rates on *β*-protein cluster size
- Details on the free energy profiles
- Additional data for the comparison with experimental results
- Lipid solubility in simulations and experiments

### Pair-wise interactions between membrane lipids and proteins

**FIG. A1.**
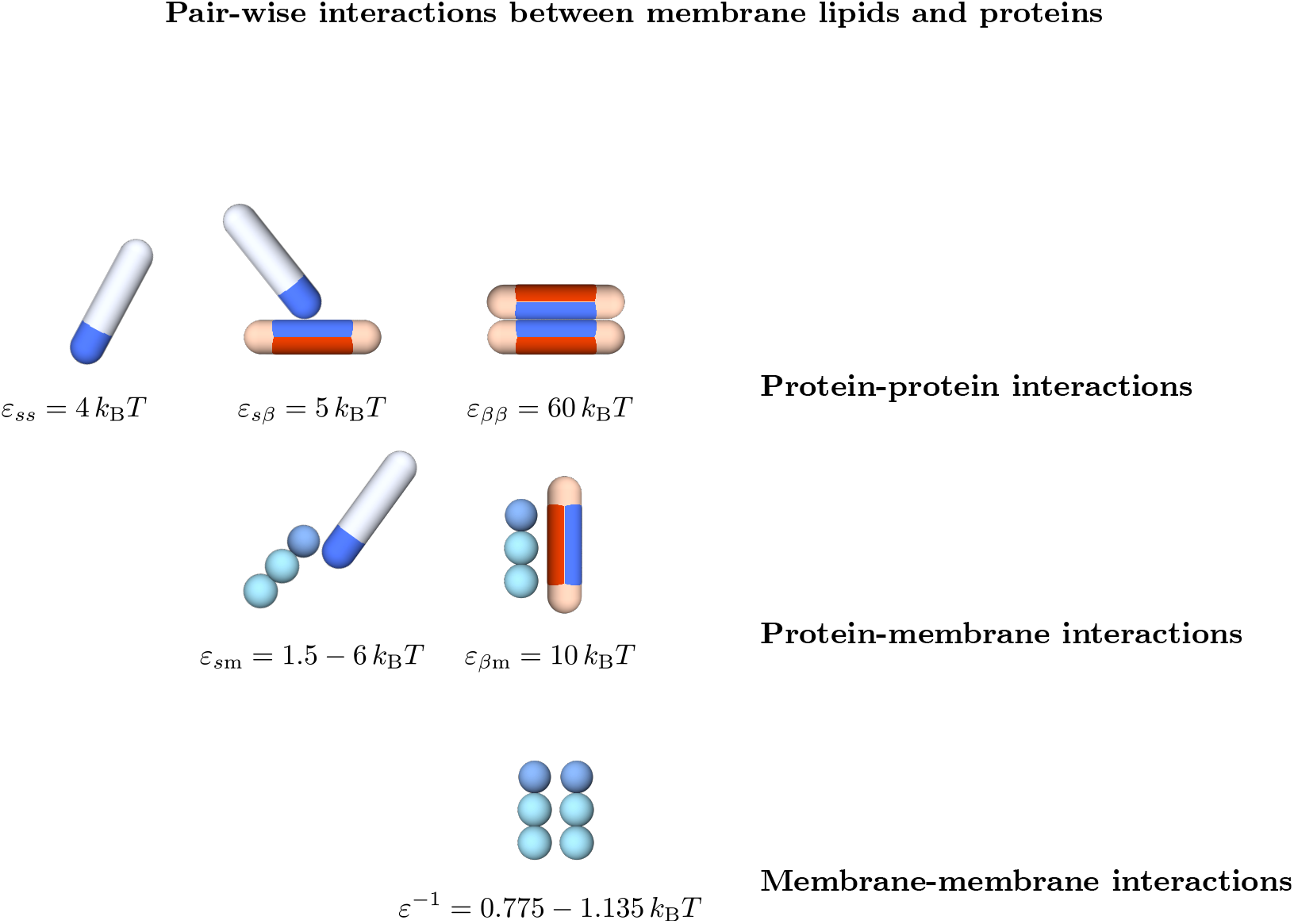
Overview of interactions in the simulation model: All possible pair interactions between proteins and membranes implemented in the simulation model.

### Remark on nucleation in solution vs membrane-assisted nucleation

We have confirmed that for a conversion free energy barrier of Δ*F*_s→*β*_ = 20 *k*_B_*T* nucleation in solution never occurs within our simulation time (4* 10^7^ Monte-Carlo time steps). This is the case using the protein number concentration of 0.002, a protein self-interaction *ε_ss_* = 4 *k*_B_*T* and a switching attempt frequency of 100, the same conditions used in the simulations in the presence of the lipid bilayer. Under these condition, with a *β*-membrane interaction *ε*_*β*m_ = 20*k*_B_*T* nucleation occurs on average after 1.2 * 10^5^ time steps in the presence of a fluid membrane (*k*_B_*T*/*ε* = 1.015) in intermediate the protein-membrane affinity regime at *ε*_*s*m_ = 3.5 *k*_B_*T*. Hence, membrane-assisted nucleation in our model can be significantly faster than homogeneous nucleation, as reported experimentally.

Even though our model is simple, the large number of lipids and the need for a substantial nucleation statistics prevents us from exploring slow nucleation mechanisms, which is needed to be able to understand the requirements for fast nucleation. To be able explore the slow nucleation rates realised in the case of a gel-like membrane and at both high and low values of protein-membrane affinity *ε*_*s*m_ we reduced the conversion free energy barrier to Δ*F*_s→*β*_ = 10 *k*_B_*T*. At the same time the *β*-membrane interaction was scaled proportionally to *ε*_*β*m_ = 10 *k*_B_*T*. In addition a condition was imposed such that homogeneous nucleation in solution cannot occur, which would be the case for the lower conversion barrier. This condition requires that a *s* → *β* conversion can occur only in the presence of a lipid molecule.

Imposing this scheme allows us to efficiently sample the full range of parameters in our simulation. In particular, the full range of protein-membrane affinities *ε*_*s*m_ up to saturation and the complete gel-to-fluid transition of the lipid bilayer.

### Coverage of the membrane surface by *s*-proteins

Proteins adsorb to the membrane by virtue of their tip interaction with the lipids which is controlled by the value of *ε*_*s*m_. Increasing the value of *ε*_*s*m_ leads to a Langmuir-like behaviour of the membrane coverage *θ*, as can be seen in Fig. A2(a). Note that due to the fact that the membranes are simulated at zero lateral tension, their area decreases when reducing fluidity due to closer packing of the lipids. The averaging for each value is a configurational and temporal average over 10 realisations and 20 frames separated by 20000 Monte-Carlo time steps each. It is interesting to note that the coverage of the membrane in the gel regime at *k*_B_*T*/*ε* = 0.775,0.815 is enhanced with respect to the fluid regime. In particular, we can see in Fig. A2(b) that the surface density of bound monomers is significantly higher for membranes of the lowest fluidities with respect to highly fluid membranes. This effect, which increases with higher *ε*_*s*m_-values, stems from the fact that more ordered membrane surfaces promote the binding of isolated monomers in the indented pockets of closely packed lipids. This is accompanied by the formation of a brush-like configuration of bound proteins. Figure A2(c) shows the enhancement of bound dimers on the two most gel-like membranes at intermediate membrane-protein affinities *ε*_*s*m_ between 3.5 and 4.5 *k*_B_*T*. This phenomenon also translates to the trimer density, as seen in Fig. A2(d).

We can understand the behaviour of the membrane coverage in more detail by considering the average cluster size of bound soluble proteins.

### Cluster size distribution of bound *s*-proteins

After equilibration of the Langmuir-like binding isotherm we measured the size distribution of oligomers bound to the membrane. The criterion for two proteins to be counted in the same oligomer is that they must interact through their tips while being bound to the membrane. Note that in this case the oligomer size is obtained before the proteins can switch to their *β*-sheet forming conformation.

The average cluster size is plotted in Fig. A3(a). It directly derives from the number of bound oligomers up to size 10, as shown in Fig. A2. At the lowest binding affinities the average cluster size can drop below one (only monomers) due to the fact some snapshot during temporal averaging contain no bound proteins. Furthermore, the average cluster size rises more steeply with *ε*_*s*m_ for less fluid membranes. Above *ε*_*s*m_ = 4.5 *k*_B_*T* the average cluster size for the two least fluid membranes saturates and even drop as *ε*_*s*m_ is increased further. This can be explained by the fact that the abundance of bound proteins further drives the lipids in to a close-packed state. There proteins bind strongly in the resulting dents between the close-packed lipid heads which hinders the formation of oligomers and leads to an excess in bound monomers.

Effectively, this results in a opposite scaling of the average cluster size with the fluidity above and below the threshold *ε*_*s*m_ = 4.5 *k*_B_*T*, which signals a binding regime of strong protein binding. This scaling is shown Fig. A2(b).

**FIG. A2.**
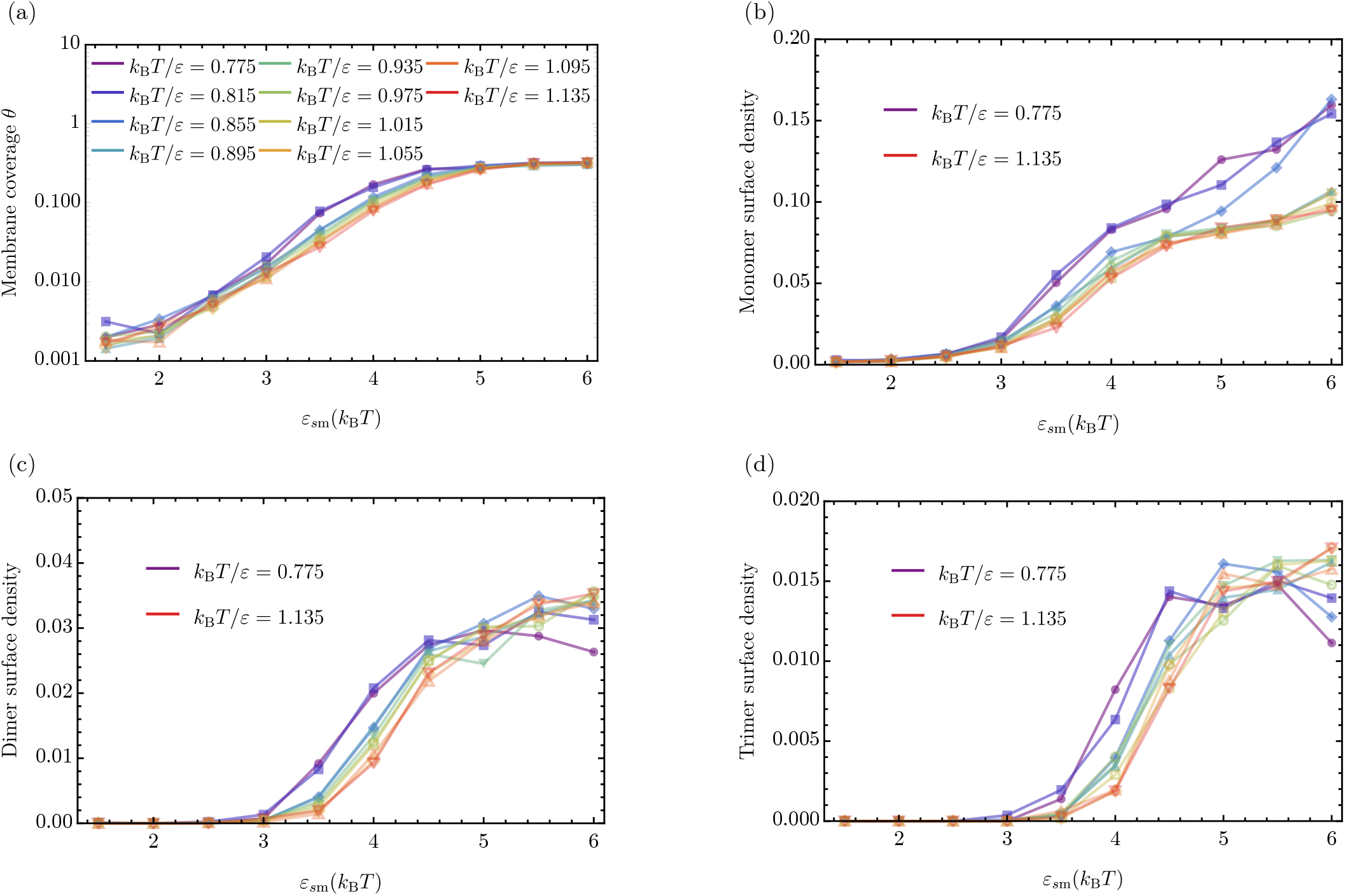
Membrane coverage characteristics of soluble proteins as a function of *ε*_*s*m_ and fluidity: **(a)** Average number of adsorbed proteins per unit area on the lipid membrane as a function of the protein-membrane fluidity. Different colours indicate ten different values of fluidity in the range *k*_B_*T*/*ε* ∈ {0.775, 0.815,&, 1.135}. Average number per unit area of oligomers of increasing size: **(b)** monomers, **(c)** dimers, and **(d)** trimers.

### Distribution of area per lipid

In order to shed light on the influence of bound protein proteins we computed the distribution of areas per lipids as a function of both the membrane-protein affinity *ε*_*s*m_ and the membrane fluidity. The area per lipid distribution was obtained by Voronoi tesselation.

Figure A4 shows the average area per lipid as a function of the fluidity. Firstly, we directly can see the signature of the gel-to-fluid membrane phase transition as a jump in the average area per lipid as *k*_B_*T*/*ε* is increased, i.e. as the strength of the hydrophobic lipid tail attraction is decreased. In addition we observe that the three cases with the highest membrane-protein affinities exhibit a shifted phase boundary between gel and fluid. The adsorption of proteins delays the gel-fluid transition towards higher *k*_B_*T*/*ε*-values. Or put differently, the adsorption of *s*-proteins to the membrane can induce a fluid-to-gel transition.

### Cluster size distribution of bound *β*-proteins

In addition to the case of soluble proteins, we also recorded the oligomer size distribution of converted *β*-proteins. Simulations were run until 20 proteins have converted to their *β*-state, then the cluster size distribution was recorded. Here, the criterion for being counted in the same cluster is that two proteins have to be interacting through their side patches in the *β*-conformation. The results can be seen in Fig. A5.

The average cluster size of *β*-prone prone proteins depends on both the membrane-protein affinity *ε*_*s*m_ and the membrane fluidity. High membrane-protein affinities entail a high membrane coverage, which in turn leads to the quick elongation of nucleated *β*-proteins. This manifests itself in a average cluster size of 20 for the lowest fluidity above affinities of *ε*_*s*m_ = 4.5 *k*_B_*T*.

**FIG. A3.**
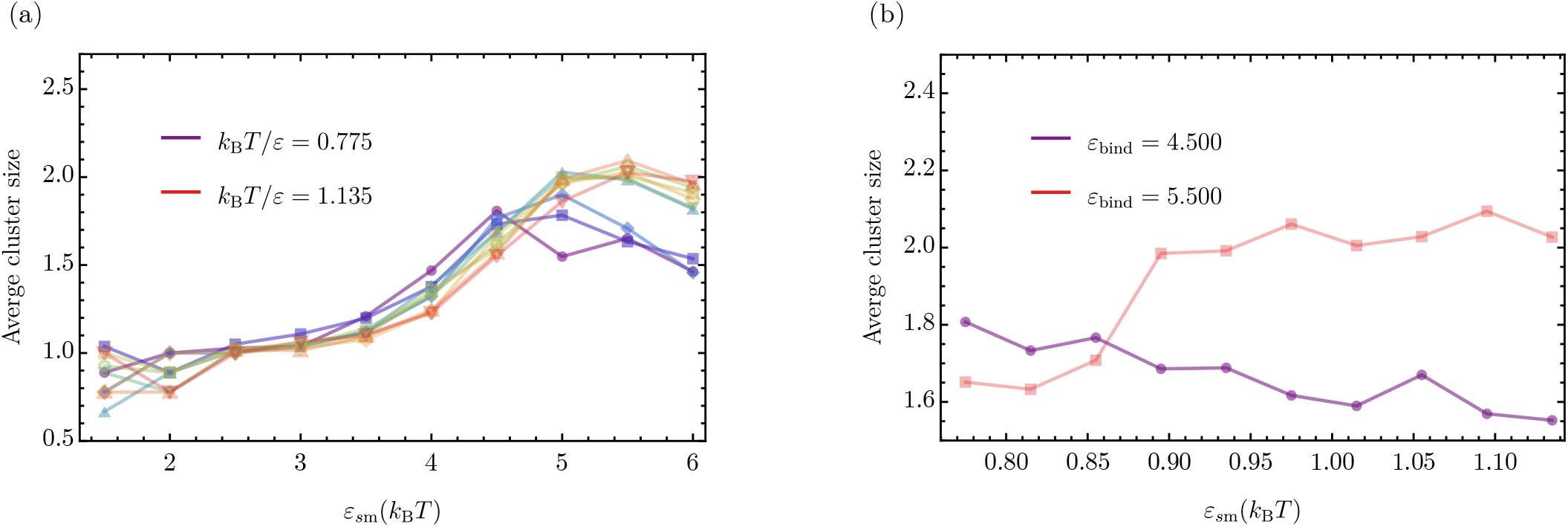
Average cluster size of bound protein oligomers. **(a)** Average cluster size of soluble protein oligomers bound to the lipid bilayer. Different colours indicate ten different values of fluidity *k*_B_*T*/*ε* ∈ {0.775, 0.815,&, 1.135}. **(b)** Slice through **(a)** at the constant protein-membrane affinities *ε*_*s*m_ = 4.5kBT and *ε*_*s*m_ = 5.5*k*_B_*T* showing an increase (decrease) of the average oligomer size with growing membrane fluidity.

**FIG. A4.**
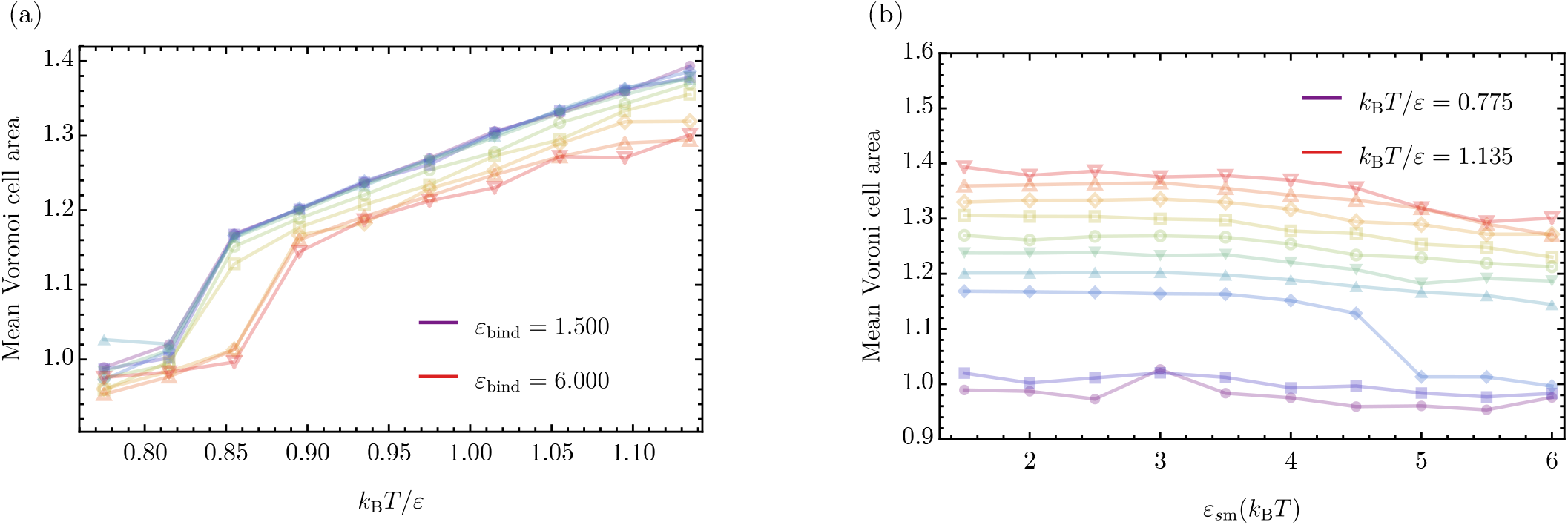
Average area per lipid: Average area per lipid as a function of **(a)** membrane fluidity and **(b)** membrane-protein affinity obtained via Voronoi tesselation.

We can observe that increasing fluidity tends to reduce the average *β*-cluster size due to the increased tendency towards producing mixed lipid-protein clusters.

### Dependence of the nucleation rates on the cluster size

As discussed in the main text, the nucleation rates in Fig. 3 are determined by the time step where the first *β*-prone dimer is formed. In Fig. A6 we report the dependence of the nucleation rate of clusters on the size of the cluster. The main conclusion we can draw from this analysis is that the formation of higher order *β*-clusters is suppressed in the regime of small membrane-protein affinities and high fluidities. This can be explained by the consideration that fast formation of higher order *β*-cluster on the one hand requires a local environment enriched in bound proteins, which is not provided at low *ε*_*s*m_-values and on the other hand can be inhibited by the absorption of *β*-proteins into the membrane core facilitated at high fluidities.

**FIG. A5.**
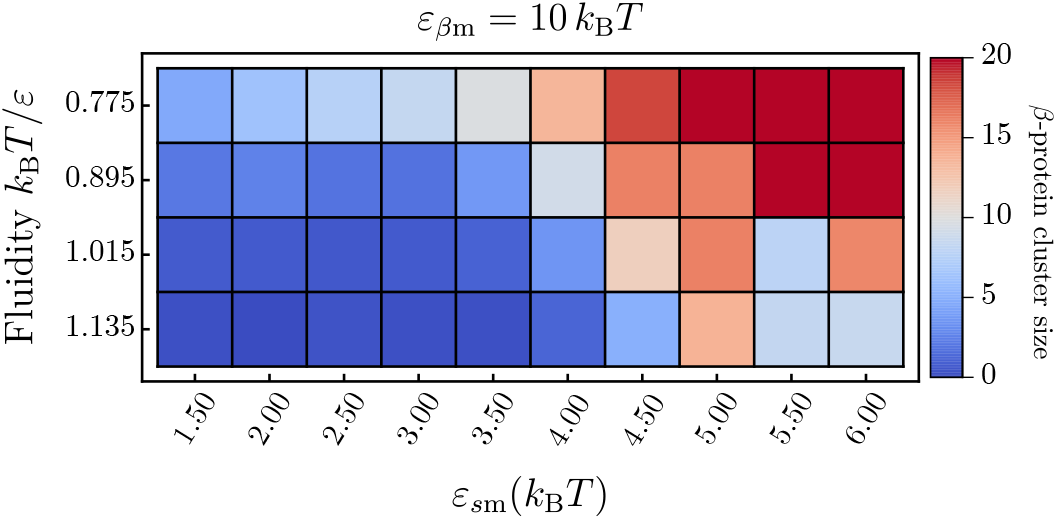
Average size of *β*-protein clusters: Average cluster size distribution once 20 *β*-prone proteins were formed.

**FIG. A6.**
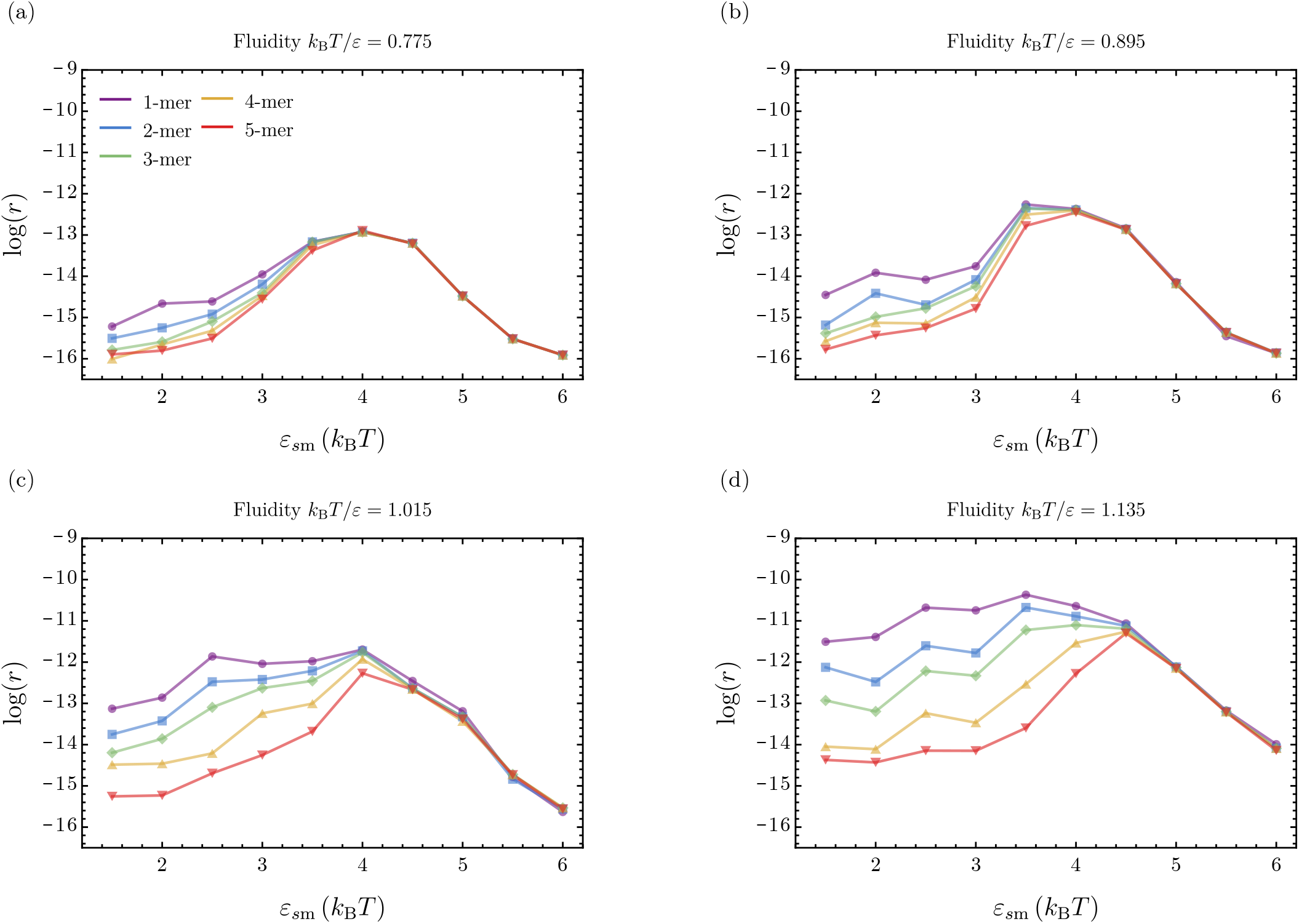
Nucleation rate for different cluster sizes. The four panels show the nucleation rates as a function of ε_sm_ for the four different fluidities studied in this work. Each panel reports the rate for forming a *β*-monomer up to a *β*-pentamer as the membrane fluidity *k*_B_*T*/*ε* and the membrane-protein affinity *ε*_*s*m_ are varied.

### Details on the free energy profiles

The potentials of mean force (PMF) for both protein species were obtained using Umbrella simulations in which one isolated protein was brought into interaction with the lipid bilayer by lowering the protein in a stepwise fashion from solution into the the centre of the lipid membrane.

Therefore, these simulations inherently cannot account for cooperative effects between proteins but will only reflect the influence of the membrane on the conversion probability of a single isolated protein.

**FIG. A7.**
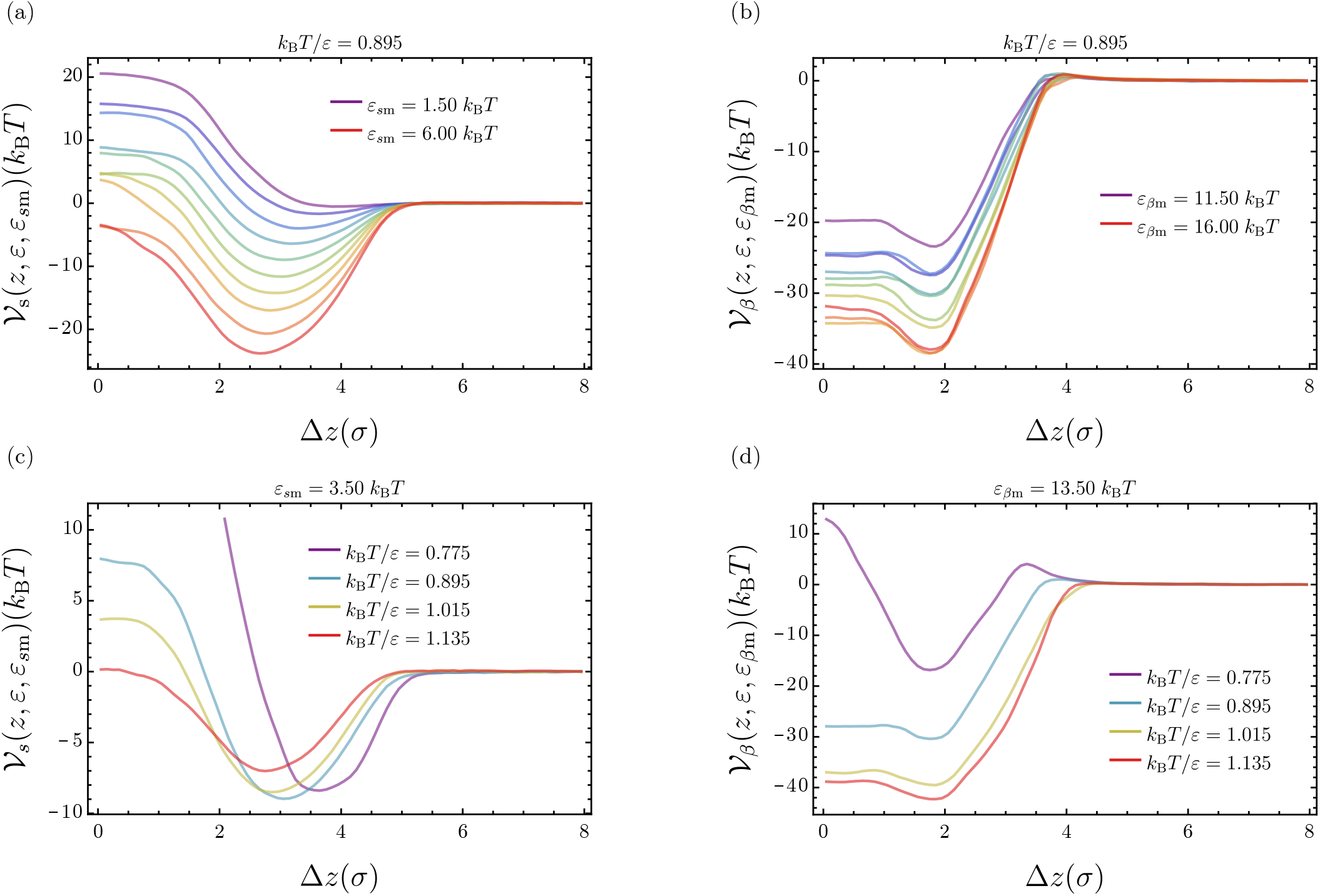
Free energy profiles of the soluble and *β*-like protein. The four panels show a selection of free energy profiles: **(a)**, **(b)** at fixed fluidity and **(c)**, **(d)** at fixed affinity. **(a)** Free energy profile of the soluble protein at fixed fluidity *k*_B_*T*/*ε* = 0.895 and varying affinities *ε*_*s*m_. **(b)** Free energy profile of the *β*-like protein at fixed fluidity *k*_B_*T*/*ε* = 0.895 and varying affinities *ε*_*β*m_. **(c)** Free energy profile of the soluble protein at fixed affinity *ε*_*s*m_ = 3.50*k*_B_*T* and varying fluidity. **(d)** Free energy profile of the *β*-like protein at fixed affinity *ε*_*β*m_ = 13.50kBT and varying fluidity.

From the umbrella simulation we obtained the two potentials of mean force for the soluble and *β*-like protein, denoted by 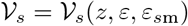 and 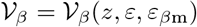, respectively. Here, the variable *z* designates the centre-of-mass separation between the membrane and the protein.

A single protein is placed at a sufficient distance from the membrane to be out of interaction range. Then the protein is inserted into the membrane using a biasing potential, as illustrated in Fig. 4. Subsequently, the weighted histogram analysis method [35] is used to compute 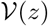. The resulting PMFs reflect the corresponding free energy of binding in the protein-membrane system.

We distinguish the free energy profiles for the *s*- and *β*-states of the protein that are denoted by 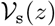 and 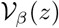, respectively. The minimum position zmin,s reflects the average equilibrium binding position of the protein along the membrane normal.

#### Variation with protein-membrane affinity

As the respective affinities are increased, the depth of both the PMFs of the soluble and *β*-like proteins increase reflecting the stronger binding of the protein to the membrane, as can be seen in Fig. A7(a) and (b) for *k*_B_*T*/*ε* = 0.895. In addition to this, we observe that the minimum of 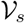 shifts also in position to lower values of the centre-of-mass separation, which is not the case for 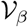. This results from the model assumption that soluble proteins can also interact with the two hydrophobic lipid tails.

**FIG. A8.**
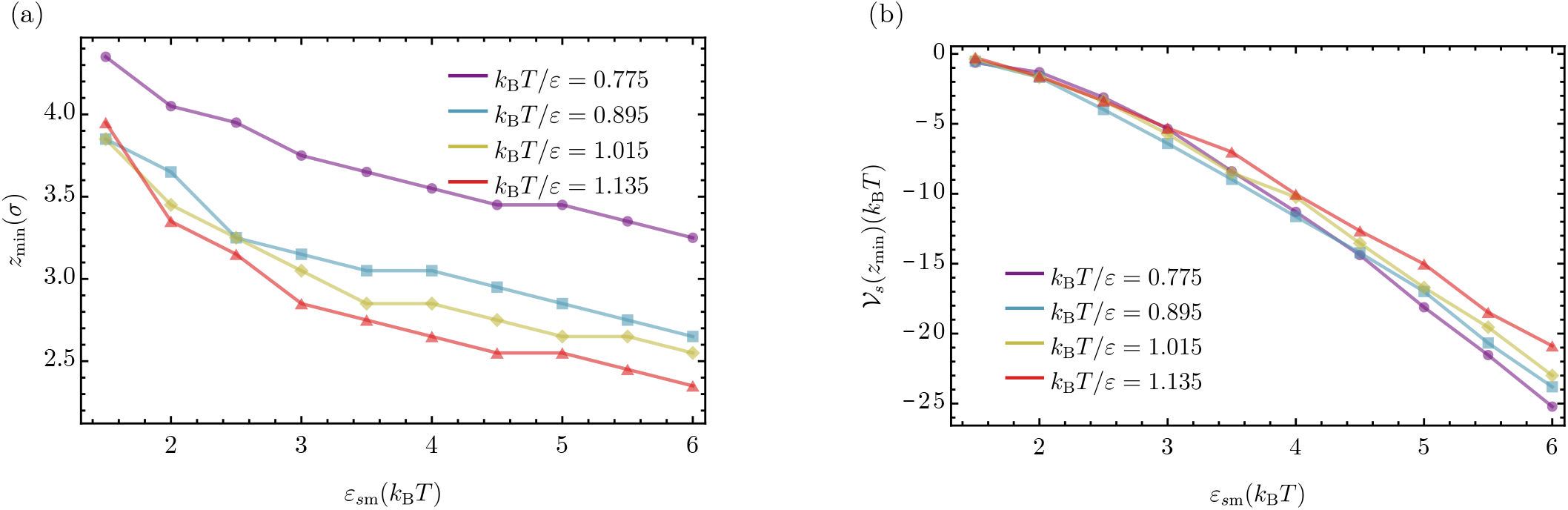
Minima of the soluble potential of mean force: (a) Position of the minima of 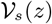 as a function of ε_sm_. Different colours indicate the four different membrane fluidities discussed in this work. (b) Depth of the free energy profile 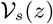 at the minimum position *z*_min_ as a function of membrane fluidity and membrane-protein affinity.

At the same time the density profile of the membrane lipids is not affected by increasing the affinity. From this we can conclude that the soluble protein insert further into the lipid bilayer as its affinity to the membrane is increased, which is a result of the specific lipid interaction profile we have chosen.

#### Variation with membrane fluidity

Varying the membrane fluidity, the soluble PMF reacts differently. This situation is plotted in Fig. A7 (c). At the lowest fluidity *k*_B_*T*/*ε* = 0.775, the protein experiences a strong repulsive branch of the PMF, while being bound in a relatively narrow well on top of the lipid membrane. In the more fluid phases the resulting PMF is markedly softer. We observe both a decrease in the depth of the PMF and and shift to lower values of Δ*z*_cm_. However, in contrast to the constant fluidity case, now the density profile of the membrane is changed as well which results from the thinning of the membrane as its phase state changes from gel to fluid. The average *z* position of the lipid head in the top leaf of the membrane decreases from 2.4*σ* ± 0.35*σ* to 1.92*σ* ± 0.64*σ* indicating that the membrane thins and at the same time the lipid head experiences higher fluctuations in *z*. Going across the gel-fluid transition, the minima of the soluble PMF shifts by more than *σ* which indicates that the shift in the PMF position is a combination of the thinning of the membrane and a higher mobility along the membrane normal due to higher fluidity.

A slightly different situation arises for the *β*-like protein as shown in Fig. A7(d). If the membrane is in its gel phase, the PMF 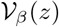 exhibits a barrier since the *β*-proteins do not directly bind to the hydrophilic lipid heads. As the fluidity of the membrane is increased, this barrier vanishes and the resulting value of the PMF minimum decreases with higher fluidities.

In summary, the membrane fluidity in our simulation model has a direct effect on the characteristics of the membrane-protein interaction. As can be seen in Fig. A8(a) both the membrane fluidity and the protein-membrane interaction have a significant effect on the position of the PMF minima for the soluble protein. The resulting affinity is shown in Fig. A8(b). We see that at high affinities, the protein is more strongly bound at low fluidities than at high fluidities. This results from the closer packing of the lipid head, and bound protein can arrange in the resulting lattice-like surface of the membrane surface maximising its binding energy.

## COMPARISON OF SIMULATION RESULTS WITH EXPERIMENTAL DATA

In order to allow for a quantitative comparison to available experimental data we consider the variation of amyloid nucleation rates as a function of the average area per lipid. Specifically, we consider the relative increase in rates and area per lipid between different fluid membranes with respect to a gel membrane given by *r*_fluid_/*r*_gel_ and *A*_fluid_/*A*_gel_, respectively.

Plotting *r*_fluid_/*r*_gel_ against *A*_fluid_/*A*_gel_ for the two cases discussed in the main text (Fig. 3), we can observe a markedly different scaling. If *ε*_*β*m_ = 0 *k*_B_*T*, *r*_fluid_/*r*_gel_ shows a very weak dependence on the increase in area per lipid, which is cause by increasing membrane fluidity. In contrast to this the interacting case *ε*_*β*m_ = 0 *k*_B_*T* exhibits a significantly higher variation with the increase in area per lipid caused by the increased exposure of hydrophobic content through membrane defects as fluidity grows. To quantify the different scalings we fitted the two data sets which provides us with two exponents *λ* for the fitting function of the form

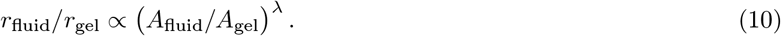

The fits result in *λ* = 0.42 for the case *ε*_*β*m_ = 0 *k*_B_*T* and *λ* = 14.2 for the case *ε*_*β*m_ = 10 *k*_B_*T*, as illustrated in Fig. A9.

**FIG. A9.**
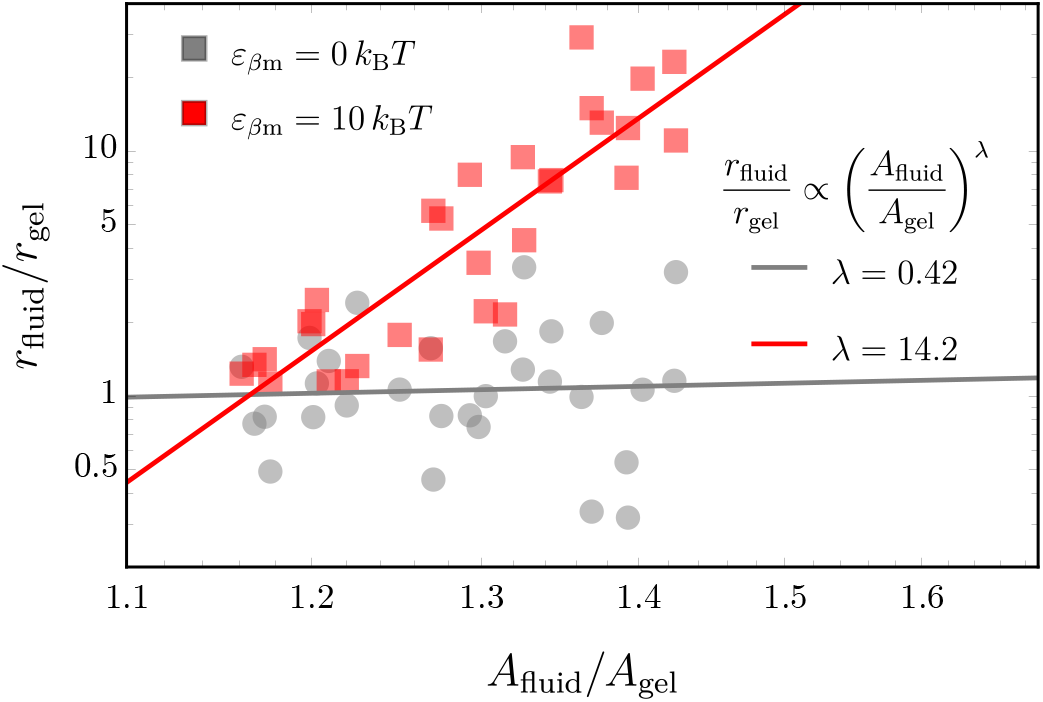
Increase in nucleation rates as a function of area per lipid: The relative increase in nucleation rates is plotted against the relative increase in area per lipid for the two cases with and without interactions between the lipid tails in the *β*-prone proteins. In addition, fits to the data points are shown to retrieve the scaling exponent *λ*.

## LIPID SOLUBILITY IN SIMULATIONS AND EXPERIMENTS

Experiments in Ref. [16] have shown that decreases the length of the acyl chain of saturated lipids from (16: 0)_2_ (DPPS) to (14: 0)_2_ (DMPS) and (12: 0)_2_ (DLPS) increases to solubility (or free energy of transfer) Δ*G* from approximately –55*kJ*mol^−1^ to –48*kJ*mol^−1^ and –41*kJ*mol^−1^, respectively. Under the experimental condition in this Reference, the (16: 0)_2_ lipid vesicles are in the gel phase, whereas the other two are in the fluid phase.

The corresponding values of the lipid free energy of transfer in our simulations exhibit the same trend when reducing the interaction strengths between the lipid tails (i.e. going from the gel to the lipid phase). However, a precise match of the numerical values of the lipid solubility cannot be achieved from such a general lipid membrane model.

For the different fluidities *k*_B_*T*/*ε* = 0.775,0.895,1.015 and 1.135 the value of the lipid solubility in simulations is approximately Δ*G* = –45*kJ*mol^−1^, –35*kJ*mol^−1^, –25*kJ*mol^−1^ and –15*kJ*mol^−1^, respectively.

## REFERENCES

[1] C. M. Dobson, Cold Spring Harbor Perspectives in Biology 9, a023648 (2017).

[2] T. P. Knowles and R. Mezzenga, Advanced materials (Deerfield Beach, Fla.) 28, 6546 (2016).

[3] S. Auer, A. Trovato, and M. Vendruscolo, PLoS Computational Biology 5, e1000458 (2009).

[4] C. Cabaleiro-Lago, O. Szczepankiewicz, and S. Linse, Langmuir 28, 1852 (2012).

[5] T. John, A. Gladytz, C. Kubeil, L. L. Martin, H. J. Risselada, and B. Abel, Nanoscale 10, 20894 (2018).

[6] A. Morriss-Andrews and J.-E. Shea, The Journal of Chemical Physics 136, 065103 (2012).

[7] R. Vácha, S. Linse, and M. Lund, Journal of the American Chemical Society 136, 11776 (2014).

[8] M. Törnquist, T. C. T. Michaels, K. Sanagavarapu, X. Yang, G. Meisl, S. I. A. Cohen, T. P. J. Knowles, and S. Linse, Chemical Communications 54, 8667 (2018).

[9] A. Šaric, A. K. Buell, G. Meisl, T. C. Michaels, C. M. Dobson, S. Linse, T. P. Knowles, and D. Frenkel, Nature Physics 12, 874 (2016).

[10] P. K. Auluck, G. Caraveo, and S. Lindquist, Annual Review of Cell and Developmental Biology 26, 211 (2010).

[11] G. Fusco, A. De Simone, P. Arosio, M. Vendruscolo, G. Veglia, and C. M. Dobson, Scientific Reports 6, 1 (2016).

[12] A. Khondker, R. J. Alsop, and M. C. Rheinstädter, Membranes 7 (2017), 10.3390/membranes7030049.

[13] K. A. Burke, E. A. Yates, and J. Legleiter, Frontiers in Neurology 4, 1 (2013).

[14] L. Giehm, C. L. P. Oliveira, G. Christiansen, J. S. Pedersen, and D. E. Otzen, Journal of Molecular Biology (2010), 10.1016/j.jmb.2010.05.060.

[15] C. Galvagnion, A. K. Buell, G. Meisl, T. C. Michaels, M. Vendruscolo, T. P. Knowles, and C. M. Dobson, Nature Chemical Biology 11, 229 (2015).

[16] C. Galvagnion, J. W. P. Brown, M. M. Ouberai, P. Flag-meier, M. Vendruscolo, A. K. Buell, E. Sparr, and C. M. Dobson, Proceedings of the National Academy of Sciences 113, 7065 (2016).

[17] M. Yang, K. Wang, J. Lin, L. Wang, F. Wei, J. Zhu, W. Zheng, and L. Shen, Langmuir 34, 8408 (2018).

[18] A. Choucair, M. Chakrapani, B. Chakravarthy, J. Katsaras, and L. J. Johnston, Biochimica et Biophysica Acta - Biomembranes 1768, 146 (2007).

[19] M. Bokvist, F. Lindström, A. Watts, and G. Gröbner, Journal of Molecular Biology 335, 1039 (2004).

[20] F. Hane, E. Drolle, R. Gaikwad, E. Faught, and Z. Leonenko, Journal of Alzheimer’s Disease 26, 485 (2011).

[21] J. J. Kremer, M. M. Pallitto, D. J. Sklansky, and R. M. Murphy, Biochemistry (2000), 10.1021/bi0001980.

[22] J. Habchi, S. Chia, C. Galvagnion, T. C. Michaels, M. M. Bellaiche, F. S. Ruggeri, M. Sanguanini, I. Idini, J. R. Kumita, E. Sparr, S. Linse, C. M. Dobson, T. P. Knowles, and M. Vendruscolo, Nature Chemistry 10, 673 (2018).

[23] E. I. O’Leary, Z. Jiang, M. P. Strub, and J. C. Lee, Journal of Biological Chemistry 293, 11195 (2018).

[24] I. R. Cooke and M. Deserno, Journal of Chemical Physics 123 (2005), 10.1063/1.2135785, arXiv:0509218 [condmat].

[25] A. Šarić, Y. C. Chebaro, T. P. J. Knowles, and D. Frenkel, Proceedings of the National Academy of Sciences 111, 17869 (2014), arXiv:1412.0897.

[26] A. Šarić, T. C. T. Michaels, A. Zaccone, T. P. J. Knowles, and D. Frenkel, 211926 (2016), 10.1063/1.4965040, arXiv:1610.02320.

[27] G. Fusco, A. De Simone, T. Gopinath, V. Vostrikov, M. Vendruscolo, C. M. Dobson, and G. Veglia, Nature Communications 5, 1 (2014).

[28] J. A. Hebda and A. D. Miranker, Annual Review of Biophysics 38, 125 (2009).

[29] E. Terzi, G. Hölzemann, and J. Seelig, Journal of Molecular Biology 252, 633 (1995).

[30] K. Matsuzaki, Biochimica et Biophysica Acta-Biomembranes 1768, 1935 (2007).

[31] C. Galvagnion, A. K. Buell, G. Meisl, T. C. Michaels, M. Vendruscolo, T. P. Knowles, and C. M. Dobson, Nature Chemical Biology 11, 229 (2015).

[32] H. Chaudhary, V. Subramaniam, and M. M. Claessens, ChemPhysChem 18, 1620 (2017).

[33] G. Fusco, S. W. Chen, P. T. Williamson, R. Cascella, M. Perni, J. A. Jarvis, C. Cecchi, M. Vendruscolo, F. Chiti, N. Cremades, L. Ying, C. M. Dobson, and A. De Simone, Science 358, 1440 (2017).

[34] G. Fusco, S. W. Chen, P. T. F. Williamson, R. Cascella, M. Perni, J. A. Jarvis, C. Cecchi, M. Vendruscolo, F. Chiti, N. Cremades, L. Ying, C. M. Dobson, and A. D. Simone, Science 1440, 1 (2017).

[35] J. Kästner, Wiley Interdisciplinary Reviews: Computational Molecular Science 1, 932 (2011).

